# Why didn’t the nudibranch cross the ocean? Understanding biogeographic and evolutionary relationships of *Hermissenda* (Nudibranchia: Myrrhinidae) Bergh, 1878

**DOI:** 10.1101/2024.06.23.600287

**Authors:** Katherine O. Montana, Terrence M. Gosliner, Sarah C. Crews, Lynn J. Bonomo, James T. Carlton, Rebecca F. Johnson

## Abstract

In the aftermath of the 2011 east Japanese earthquake and tsunami, anthropogenic debris from the east coast of Japan floated across the Pacific Ocean to the west coast of North America. One such vessel from Iwate Prefecture arrived on the coast of Oregon, and the fouling community included specimens identified as the nudibranch *Hermissenda crassicornis*, which was previously thought to range from Japan to Baja California but has since been split into three species: *H. crassicornis* (Alaska to southern CA), *H. opalescens* (British Columbia to Baja California), and *H. emurai* (Japan, Korea, Russian Far East). Previous work suggested that all of the motile invertebrates found in the tsunami debris fouling community were either pelagic or Japanese in origin. Our study sought to determine whether the nudibranch specimens collected from the Iwate vessel were, according to the new classification system, only *H. emurai* or whether the Eastern Pacific *Hermissenda* were present as well. Results from DNA sequencing and morphological analysis suggest that specimens of *H. crassicornis*, as it is currently recognized, and *H. opalescens* were found on the vessel. **This finding indicates either that these species settled after arrival to the west coast of North America or that *H. crassicornis* and *H. opalescens* is found in Japan, suggesting *Hermissenda* ranges need to be investigated further.** Occurrence data shared on the iNaturalist platform were also used to assess current ranges. Our phylogenetic tree and haplotype network constructed from COI data from all *Hermissenda* species indicate that *H. opalescens* and *H. emurai* are most closely related with *H. opalescens* sister to the clade that contains *H. opalescens* and *H. emurai*. This study demonstrates the power of combining volunteer naturalist data with lab-collected data to understand evolutionary relationships, species ranges, and biogeography.

## Introduction

The biogeography and evolutionary patterns of nudibranchs are subject to the forces of both nature and humans. Nudibranchs are mollusks of the class Gastropoda that are prevalent in virtually every marine habitat but have been most thoroughly studied in intertidal zones. They are colloquially known as sea slugs and exhibit a wide variety of striking coloration and patterns. Nudibranchs eat a variety of benthic colonial organisms, including hydroids, tunicates, sponges, and other sessile invertebrates, and they sometimes consume mobile prey, including other nudibranchs (McDonald & Nybakken, 1997; Megina & Cervera, 2003). Although nudibranchs as a group have a wide range of prey, individual nudibranch species are typically specialists, feeding only on one type of prey (McDonald & Nybakken, 1997). Nudibranchs typically have planktonic larvae, and the presence of prey signals them to settle onto the substrate and continue their development (Hadfield, 1963). With this reproductive strategy, oceanic currents may disperse nudibranchs widely via rafting. Our study focuses on the biogeography of the genus *Hermissenda* Bergh, 1878, which is found on the coasts of both the western and eastern Pacific in the northern hemisphere and was one of the taxa found on human-made debris from the 2011 east Japan earthquake and tsunami that washed up on the coast of Oregon (Carlton et al., 2017).

The genus *Hermissenda* was recently determined to be a species complex of three species rather than one (Lindsay & Valdés, 2016). *Hermissenda* has a convoluted naming and systematic history (Fig. 1). The morphology of *H. crassicornis* (Eschscholtz, 1831) is defined by white lines on their cerata, *H. opalescens* (JG Cooper, 1863) has white tips on their cerata, and the cerata of *H. emurai* (Baba, 1937) can be variable. However, cryptic speciation (Estores-Pacheco, 2020) and generally similar patterning between the species has likely contributed to confusion regarding species delineation. Lindsay and Valdés (2016) reported that the area of overlap between the two species to be between Bodega Bay and Point Reyes in California, with *H. crassicornis* extending as far north as Alaska and *H. opalescens* ranging south to the tip of Baja California. Additional observational data in the northeastern Pacific then extended the range of *H. opalescens* north to Vancouver Island (Merlo et al., 2018). An increased area of sympatry could also mean increased opportunities for hybridization, as has been shown in both marine and terrestrial organisms (Arce-Valdés and Sánchez-Guillén, 2022; Garroway et al., 2009; Hobbs et al., 2018). The possibility of hybridization between *H. crassicornis* and *H. opalescens* also may contribute to ambiguity in regard to species delineation.

**Figure 1.**
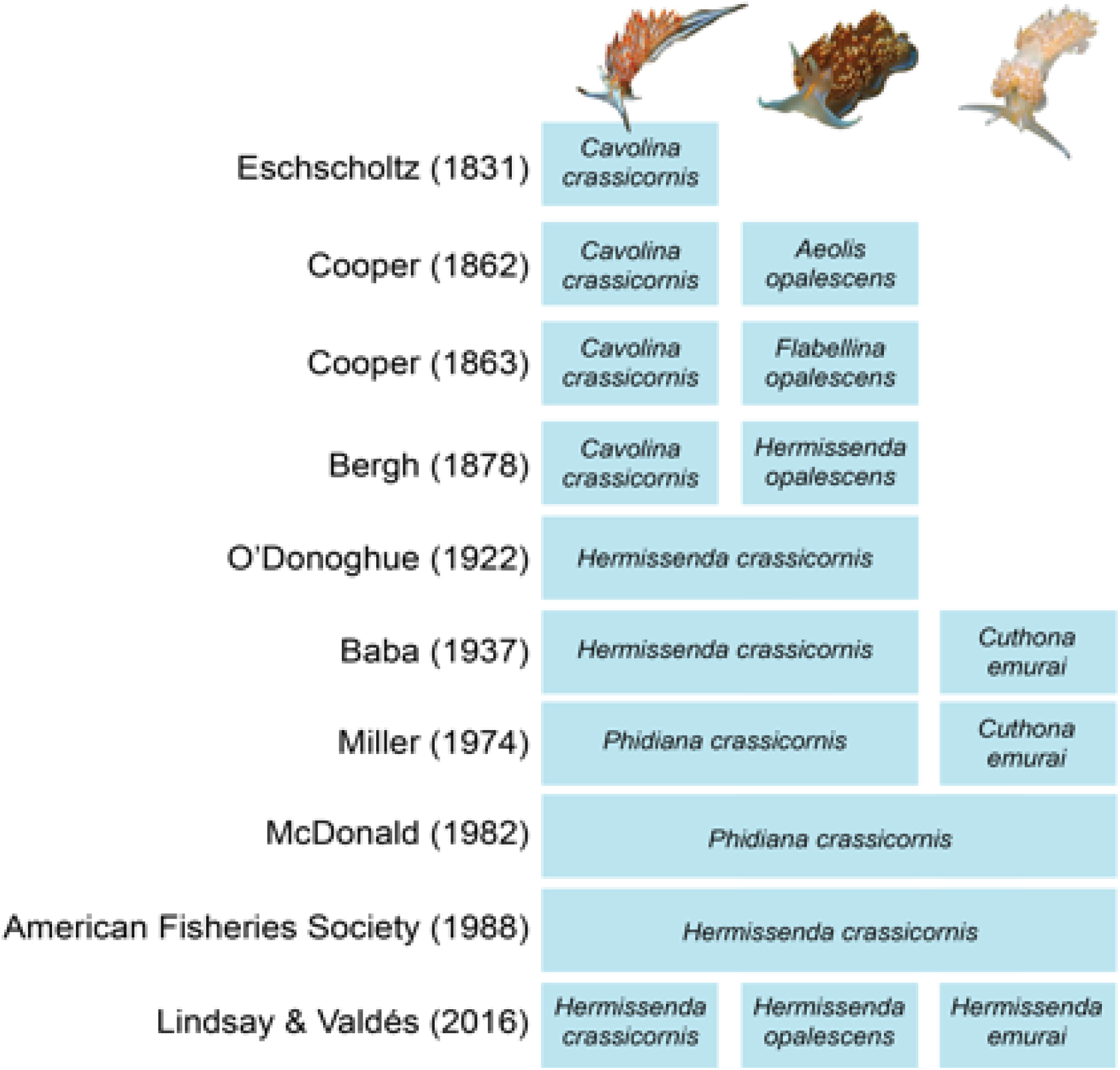
A timeline showing the taxonomic changes and citations for the family *Hermissenda* and the three species that currently comprise it. Lindsay and Valdés (2016) detailed many of the changes that are included.

Potential invasion, transoceanic dispersal, human debris, and fouling behavior are all themes in the story of the *Hermissenda* nudibranchs in our study. Invasion of marine species is often facilitated by artificial habitats and high propagule pressure (Simkanin et al., 2017). Transoceanic spread of nonindigenous marine species are common via exchange of ballast water and hull fouling (Carlton and Geller, 1993; Gollasch, 2008). Rafting, too, has previously been shown as a mechanism for species dispersal (Highsmith, 1985; Simkanin et al., 2019; Thiel and Gutow, 2005). Similar to *Hermissenda*, the cosmopolitan nudibranch *Fiona pinnata* (Eschscholtz, 1831) once thought to be a single species, has also been shown to be a species complex (Trickey et al., 2016). *Fiona pinnata* lives on macroalgal rafts, and sometimes human debris, and their fouling behavior has predisposed them to widely dispersed distribution and has likely contributed to their ability to speciate across a wide geographic area (Trickey et al., 2016). Thus, as small, fouling organisms, nudibranchs are subject to ecological processes that affect their biogeography. Our research offers a case study involving potential transoceanic dispersal via human-made rafts.

Following the March 11, 2011 east Japan earthquake and tsunami, thousands of human-made objects, including hundreds of vessels, arrived near or washed up on the shores of the northeastern Pacific Ocean and the Hawaiian Islands. Many of these objects supported marine species native to Japan (Carlton et al., 2017). One of these vessels, subsequently designated as Japanese Tsunami Marine Debris (JTMD)-BF-356, from Iwate Prefecture (see Methods) was discovered at sea in April 2015 offshore of Seal Rock, Oregon (Fig. 2). A cluster of approximately 100 nudibranchs was among the many marine organisms found in the vessel. Many of the nudibranchs were determined to be *Hermissenda crassicornis* based on the taxonomy at the time of data analysis, and others were determined to be *Dendronotus* and *Eubranchus* species (Carlton et al., 2017), genera also found in both the eastern and western Pacific (Behrens et al., 2022). Visually, the specimens resemble *Dendronotus venustus* MacFarland, 1966, and *Eubranchus rustyus* Marcus, 1961, both known only from the eastern Pacific. The identification of *Eubranchus rustyus* was confirmed by molecular data. In light of an updated classification system of *Hermissenda* that was published during the publication cycle of the Carlton et al. (2017) study (Lindsay and Valdés, 2016), we sought to confirm and possibly update the identifications of the *Hermissenda* from vessel BF-356.

**Figure 2.**
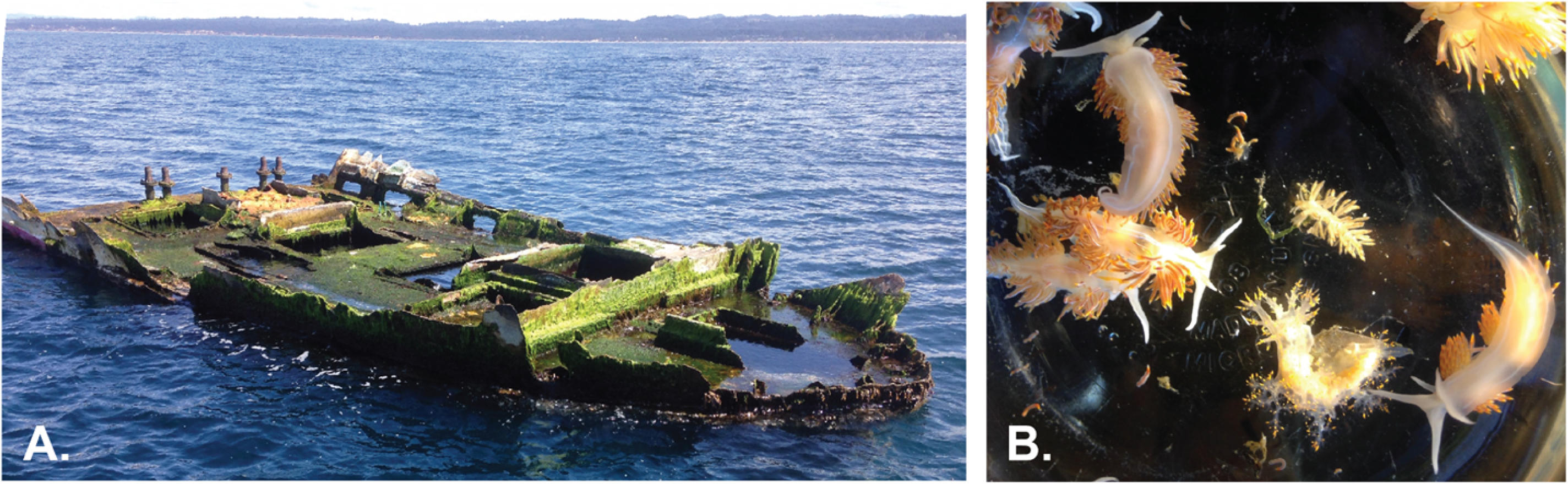
A. Vessel BF-356 discovered off the coast of Seal Rock, Oregon in April 2015, traced back to Iwate Prefecture, Japan; B. Nudibranchs found in a cluster in the vessel. Photographs by John Chapman (Carlton et al. 2018).

The Carlton et al. (2017) visual identification of *Hermissenda crassicornis* (made by G. McDonald and T. Gosliner previous to the publication of Lindsay and Valdés [2016]), was reasonable when *H. crassicornis* was thought to range across the western Pacific and northeastern Pacific. However, according to the classification system of Lindsay and Valdés (2016), if the nudibranchs had rafted from Japan, they should be re-identified as *H. emurai*. To determine the identity of the specimens from BF-356, we wanted to include all existing *Hermissenda* data on GenBank, along with other coastal Oregon specimens and the specimens found on the tsunami debris that we sequenced. Our study used molecular and morphological data to determine the identity of the nudibranchs found on vessel BF-356, leading to new understandings about the potential for nudibranch rafting of *Hermissenda* species.

## Methods

### Vessel JTMD-BF-356

On April 6, 2015, the 7.9 meter-long bow section of a large Japanese fishing vessel was found floating at sea 8 km offshore (west) of Seal Rock, Oregon, approximately 16 km south of Newport, Oregon (Carlton et al. 2017, Carlton et al. 2018, Craig et al. 2018). The vessel was identified as JTMD lost on March 11, 2011. The specific origin of the vessel in northern Honshu was likely Iwate (Craig et al. 2018). The condition of the vessel suggested that it had been on the seafloor in Japan for several years after the tsunami, and then floated free around 2013-2014; the 1- to 2-year age of Japanese fish (see below) found inside the vessel when it landed in Oregon corresponds to this estimate of time-of-departure from Japan.

The vessel was first sampled at sea on April 9, and then towed to a regional marina and extensively sampled on April 10-11, at which time the nudibranchs noted above were found in the bow internal holding tank (which would have been exposed to wash from ocean waves) (S. Rumrill, personal communication, April 2015). The live nudibranchs were then photographed by J. Chapman at the Hadfield Marine Science Center, Oregon State University, Newport, Oregon, who then provided the specimens to us.

### Molecular analyses of nudibranchs

We dissected portions of tissue for DNA extraction from the foot of 43 nudibranch specimens. Thirty-nine were from the vessel BF-356. Four *Hermissenda* were collected on our behalf by N. Treneman in 2019 from Cape Arago, Cape Sebastian, and Cape Ferrelo, OR for comparison to the species from the vessel (Table S1). Extractions were done with the Qiagen DNEasy Blood and Tissue Kit according to the manufacturer’s protocol. Cytochrome Oxidase subunit I (COI) was chosen for molecular analysis because it evolves quickly, can elucidate differences between sequences at the species level, and is compatible with existing species delineation data in *Hermissenda* and related groups (Lindsay and Valdés, 2016). The primers used for polymerase chain reaction (PCR) were LCO-1490: 5’-ggtcaacaaatcataaagatattgg-3’ and HCO-2198: 5’-taaacttcagggtgaccaaaaaatca-3’ (Folmer et al., 1994). The PCR protocols used were: an initial denaturation for 33 min at 94 °C; 40 cycles of: 94 °C at 30 s for denaturation, an annealing temperature of 46–48 °C for 30 s, and an elongation temperature of 72 °C for 45 s; and a final extension time of 7 min at 72 °C.

In addition to our lab-collected data, we included all existing *Hermissenda* COI sequences on GenBank (Table S2), as well as several closely related species as an outgroup to root our tree: *Godiva quadricolor* (Barnard, 1927), *Phyllodesmium jakobsenae* (Burghardt & Wägele, 2004), *Phidiana lascrucensis* (Bertsch & AJ Ferreira, 1974), *Nanuca sebastiani* (Er. Marcus, 1957), and *Dondice occidentalis* (Engel, 1925). Sequences were aligned using Muscle 2.8.31 (Madeira et al., 2022). Gene trees were constructed using IQTree 2 for maximum likelihood and MrBayes 3.2 for Bayesian inference (Nguyen et al., 2015; Ronquist et al., 2012). Model selection was performed by ModelFinder within the IQTree web platform (Kalyaanamoorthy et al., 2017; Trifinopoulos et al., 2016), and the maximum likelihood analysis was partitioned by codon position according to the following models: codon 1:TNe+I, codon 2: F81+F, codon 3: TN+F+G4 (Chernomor et al., 2016). The Bayesian tree was constructed using relatively equivalent models: codon 1: SYM + I, codon 2: F81, codon 3: SYM + gamma. Ten thousand ultrafast bootstraps were done in IQTree (Hoang et al., 2018), and MrBayes was run for 10 million generations, saving every 1000th generation. A haplotype network was built using PopART to illustrate the relationships in our specimens (Leigh and Bryant, 2015). We examined species delimitation using the automatic barcode gap discovery website (ABGD) under Jukes Cantor, Kimura K80, and simple distance criterion, with X=1.5, Pmin=0.001, Pmax=0.1, steps=10, and Nb bins=20 (Puillandre et al., 2012). Additionally, we ran a coalescent-based ASTRAL tree estimation to further interrogate the relationships between the specimens.

Scanning electron microscopy (SEM) was used to image the radulae to observe minute morphological differences between *Hermissenda opalescens* and *H. crassicornis* from vessel BF-356, using a single specimen of each; we determined the identity of each species once we received sequencing results. The SEM samples were coated with gold/palladium using a Cressington 108 Auto vacuum sputter coater and micrographs were taken using an Hitachi SU3500 scanning electron microscope at the California Academy of Sciences. We mapped occurrence data (including community-collected data from iNaturalist) for *Hermissenda opalescens* and *H. crassicornis* to delimit their range of overlap along the west coast of North America using the R packages sf v.1.0.9, ggplot2 v.3.4.0, tidyverse v.1.3.2 (Pebesma, 2018; Wickham, 2016; Wickham et al., 2019; GBIF)].

## Results

### JTMD-BF-356 vessel biota

Aboard vessel BF-356 were more than 20 living species of Japanese invertebrates and fish (Carlton et al. 2017; Carlton et al. 2018, Table 1; Craig et al. 2018). These included Japanese sponges, mussels, crustaceans (amphipods, isopods, and barnacles), bryozoans, the Japanese yellowtail jack *Seriola aureovittata* Temminck & Schlegel, 1845 and the barred knifejaw *Oplegnathus fasciatus* (Temminck & Schlegel 1844). Also aboard were oceanic, neustonic species acquired in transit from Japan to Oregon, including pelagic crabs (*Planes marinus*), barnacles (*Lepas* sp.), nudibranchs (*Fiona pinnata*), and bryozoans (*Jellyella tuberculata*), the latter indicating a colder-water, higher latitude route across the North Pacific.

**Table 1.** The Jukes Cantor p-distances between each species from ABGD analysis.

|  | *H. opalescens* | *H. crassicornis* | *H. emurai* |
| --- | --- | --- | --- |
| *H. opalescens* | 0–1.36% |  |  |
| *H. crassicornis* | 3.44%–5.17% | 0–0.93% |  |
| *H. emurai* | 3.42%–5.21% | 2.00%–3.40% | 0.46–1.67% |

The nudibranchs collected included, as noted earlier, taxa in the genera *Hermissenda, Dendronotus,* and *Eubranchus*. *Dendronotus* and *Eubranchus* were determined via morphology, and *Eubranchus* was confirmed *E. rustyus* by a BLAST of a COI sequence we produced that we did not include in our phylogeny. The larger *Hermissenda* specimens were approximately 2.5 cm in length.

### Molecular analyses and phylogenetic relationships of *Hermissenda*

We incorporated the 658-bp COI region of a total of 43 *Hermissenda* specimens for our phylogeny, and we included 62 COI sequences from GenBank. The topology of the gene trees for the Bayesian and fast likelihood analyses showed the same support values and relationships (Fig. 3). There was 100% support for the whole *Hermissenda* clade across all support values. Low support characterized the *H. opalescens* clade and the clade containing *H. emurai* and *H. crassicornis* sister to one another. The *H. emurai* clade showed high Bayesian support (0.99) but fared lower via sh-ALRT and UFBoot supports. The *H. crassicornis* clade showed a high SH-aLRT value (92.6) but low Bayes and UFBoot supports. Intraspecific relationships between specimens are less central to this study, and, thus, their support values are not included in Fig. 3. A coalescent-based estimation of intraspecific relationships can be found in the ASTRAL phylogeny, with relatively low (0.75) supports for the *H. opalescens* and (*H. emurai*+*H. crassicornis*) clades but high support (0.95) for the clade that contains all of *Hermissenda*. (Fig. 4). The specimens from vessel BF-356 were in the *H. crassicornis* and *H. opalescens* clades, and none were found in the *H. emurai* clade.

**Figure 3.**
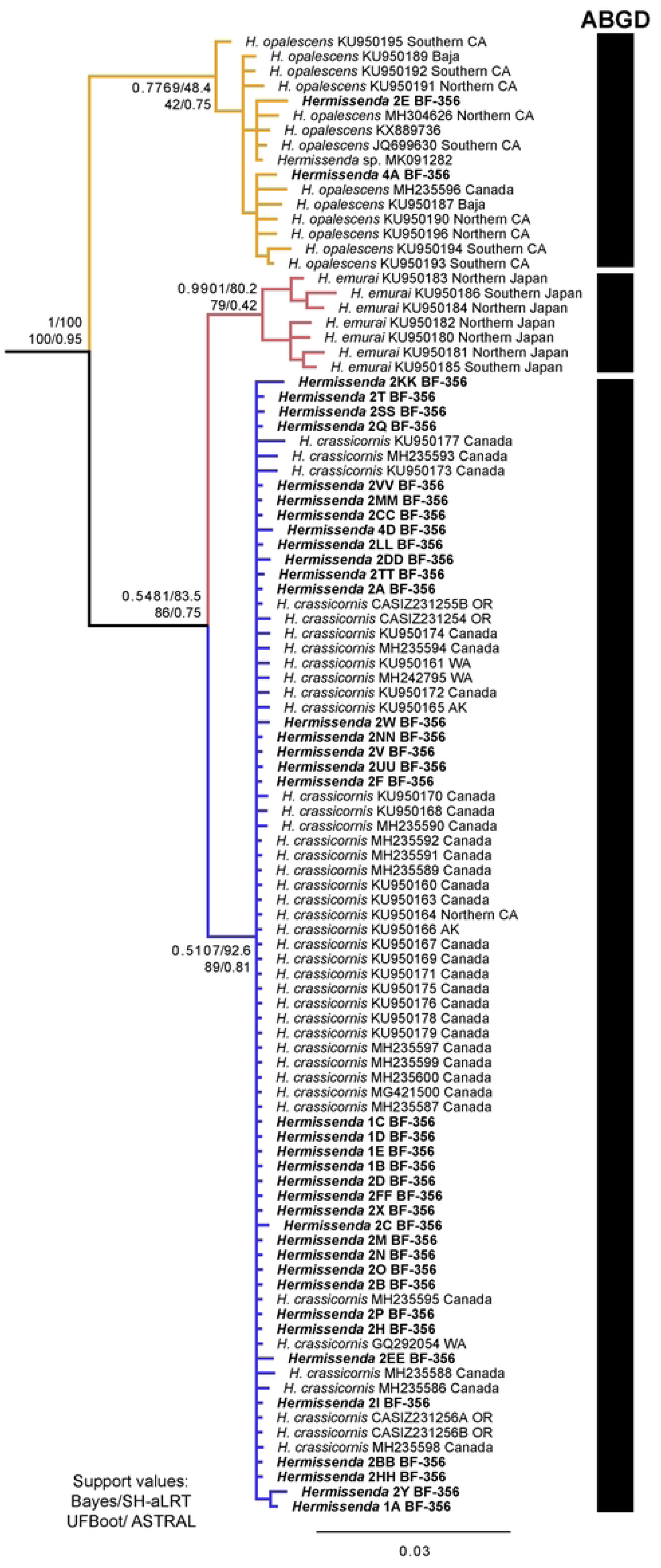
Phylogeny showing relationships of *Hermissenda* species based on COI. Support values (Bayes/SH-aLRT/UFBoot) are shown for the nodes that differentiate the species. ABGD results recovered three groups, one for each species. Samples from vessel BF-356 shown in bold. Numbers associated with BF-356 specimen names indicate the jar in which they were initially stored. Other terminal labels either have GenBank numbers and their locations, if known (state abbreviations included), or a California Academy of Sciences catalog collections number to indicate that they were new samples collected from Oregon in 2019.

**Figure 4.**
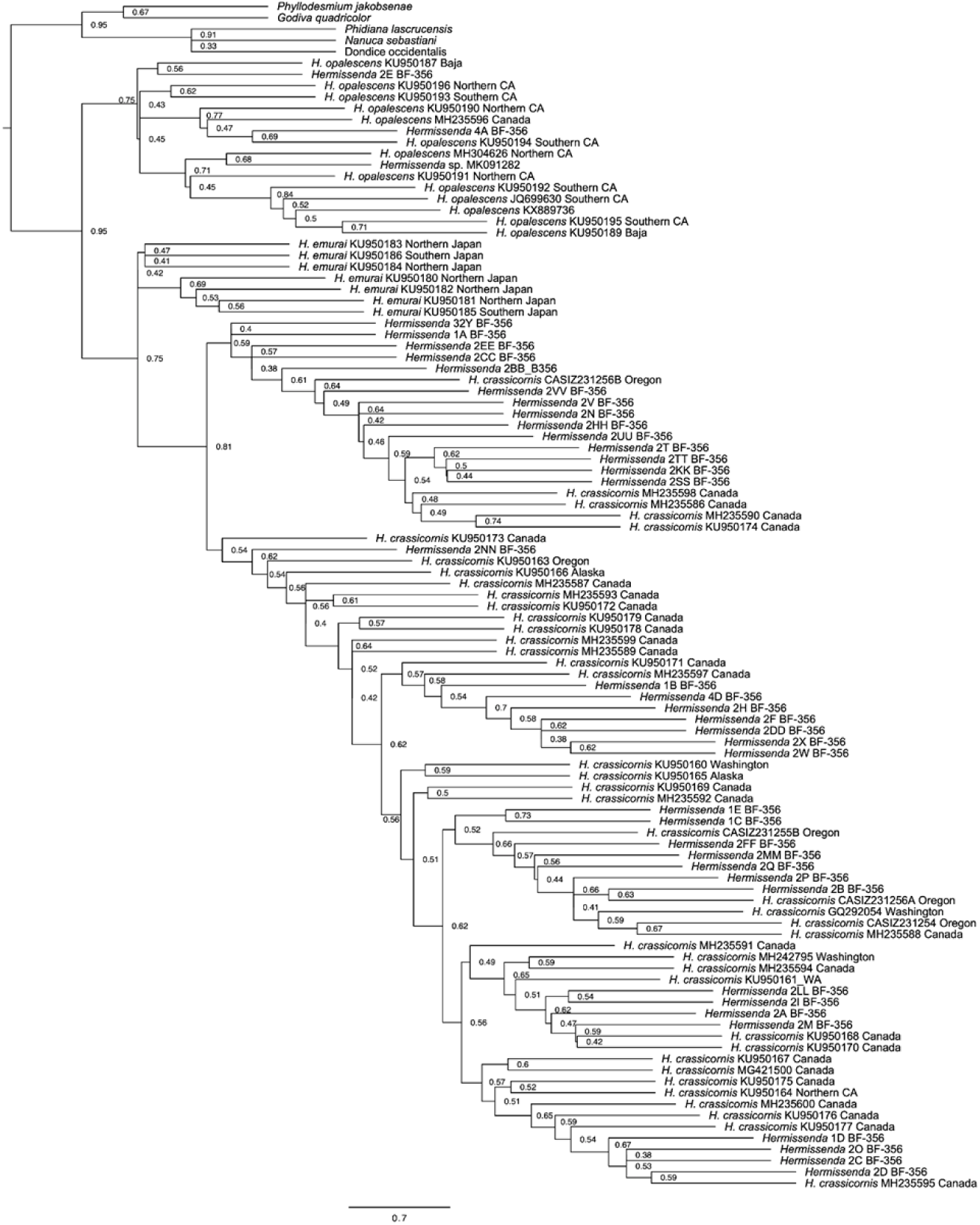
ASTRAL phylogeny showing relationships both between and within species of Hermissenda, also based on COI.

Three species groups that correspond to the three species (*H. crassicornis*, *H. emurai*, and *H. opalescens*) were recovered from the ABGD analysis (Table 1 and Fig. 3). Previous work has made haplotype networks for each *Hermissenda* species separately, but none have shown all three together (Lindsay and Valdés, 2016; Merlo et al., 2018). The haplotype network we produced shows three distinct groups, with our samples clustering according to species groups (Fig. 5). The network also shows three COI mutations present between the closest *H. crassicornis* and *H. emurai* specimens. There are 11 mutations between the closest *H. emurai* and *H. opalescens* specimens. The most distantly related *H. emurai* have seven mutations between each other. The most frequently occurring haplotype falls within the *H. crassicornis* cluster. Within the haplotype network the specimens from vessel BF-356 correspond to *Hermissenda crassicornis* and *H. opalescens* but not *H. emurai*.

**Figure 5.**
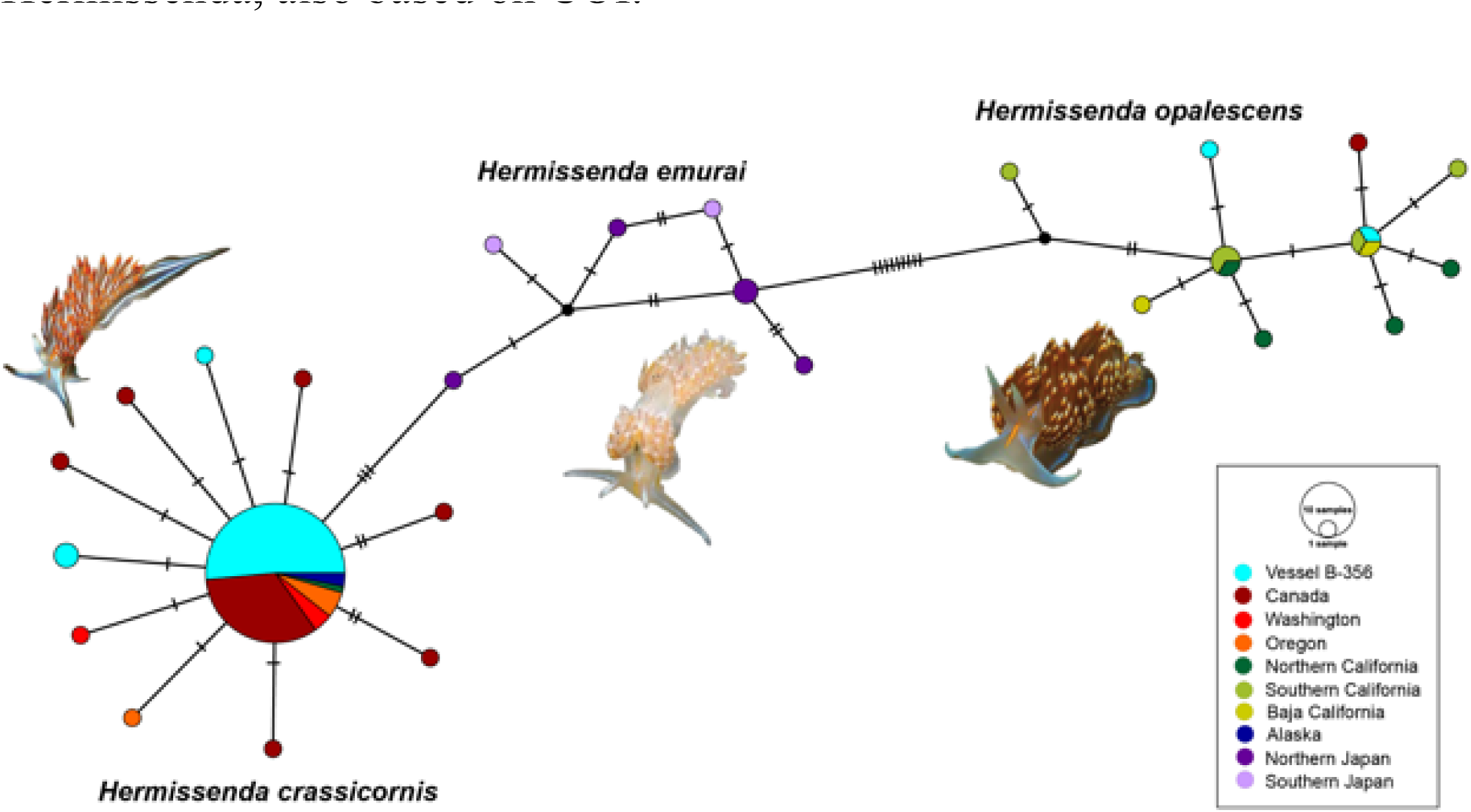
Haplotype network showing the numbers of mutations between the COI sequences and the geographic origin of each specimen.

The SEM images of the radulae of the two specimens on the BF-356 vessel showed serrations under the central cusp with denticles on either side of the cusp, typical of *Hermissenda*. *Hermissenda crassicornis* had five denticles from the central cusp, and *H. opalescens* had four denticles, with no other observable differences on the radulae (Fig. 6). While we did not have *H. emurai* specimens to examine, they are known to typically have 6–7 denticles (Lindsay and Valdés, 2016).

**Figure 6.**
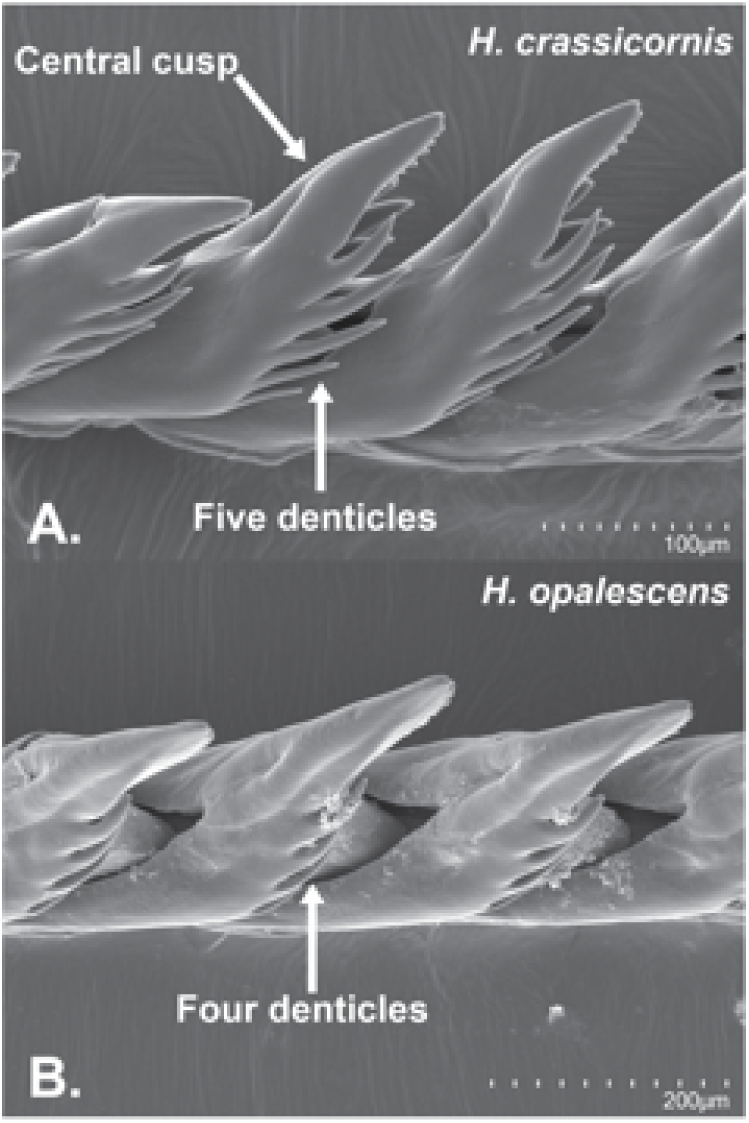
Scanning electron microscopy (SEM) images of A. *Hermissenda crassicornis* and B. *Hermissenda opalescens* radulae.

The map using GBIF data shows broad overlap between *Hermissenda crassicornis* and *H. opalescens* (Fig. 7)*. Hermissenda crassicornis* ranges from Alaska south to the southernmost coast of California. *Hermissenda opalescens* ranges from as far north as Vancouver Island south to Baja California.

**Figure 7.**
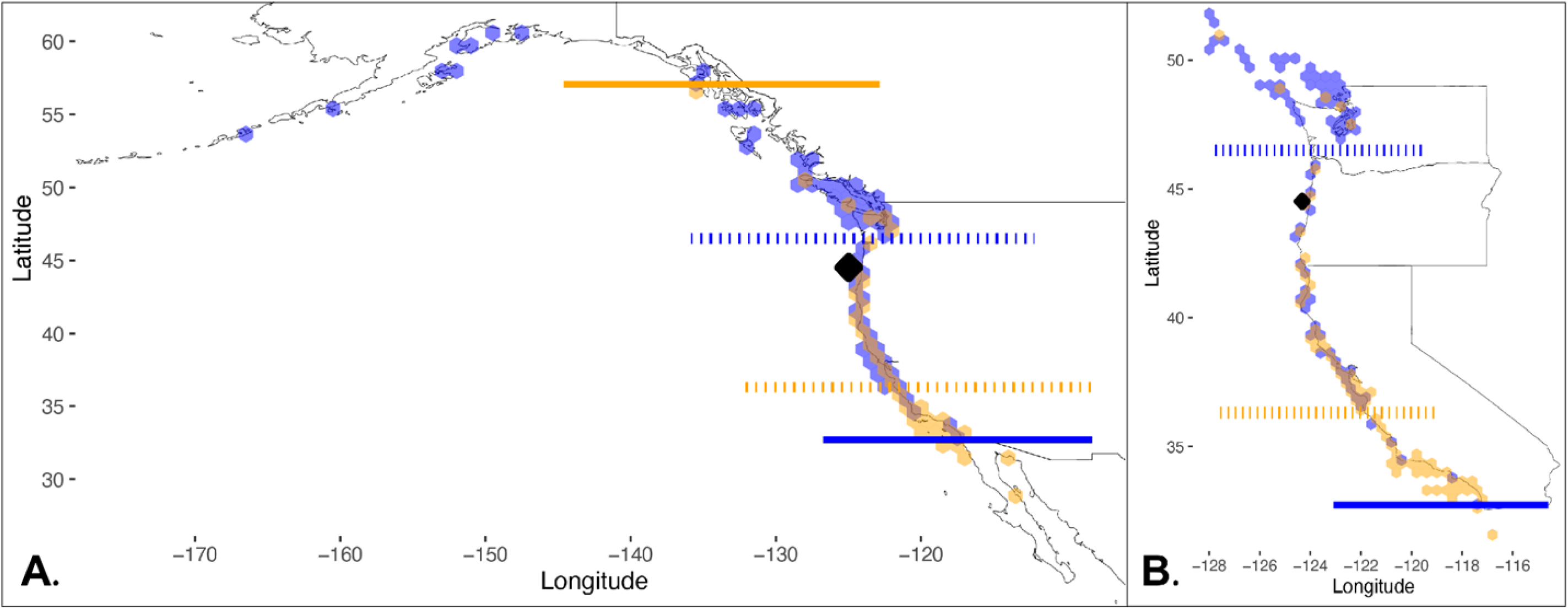
Maps made from GBIF data showing the range of overlap between *Hermissenda crassicornis* (blue) and *H. opalescens* (orange). The black diamond represents the site offshore of Seal Rock, Oregon where vessel BF-356 was found. A. Shows the entire range of both species, and B. shows a closer view of the west coast of the United States to highlight the overlap. The solid orange line represents the northernmost occurrence of *H. opalescens*, and the dotted orange line represents the mean latitude of *H. opalescens*. The solid blue line represents the southernmost occurrence of *H. crassicornis*, and the dotted blue line represents the mean latitude of *H. crassicornis*.

## Discussion

Our study revealed the identities of the nudibranchs on vessel BF-356: both *Hermissenda crassicornis* and *H. opalescens* were present, but not *H. emurai*. **This result indicates that the former two species likely did not raft from Japan following the earthquake and tsunami.** The addition of the BF-356 specimens and 2019 Oregon specimens to existing GenBank COI data for *Hermissenda* supports the currently understood species limits and relationships between *Hermissenda* species (Lindsay and Valdés, 2016). *Hermissenda crassicornis* and *H. emurai* are sister species, with both sister to *H. opalescens* (Fig. 3). Though the support values on our phylogeny were not high, the haplotype network is another line of evidence showing *H. emurai* and *H. crassicornis* as sister to one another and more distantly related to *H. opalescens*. However, we do note that even within each group, there is considerable genetic variation, with the most distantly related *H. emurai* being seven mutations away from each other. Additional variation within species can be seen in the ASTRAL phylogeny (Fig. 4). The ABGD test we ran also supported the existence of three species groups that correspond to *H. crassicornis*, *H. opalescens*, and *H. emurai* (Table 1 and Fig. 3). Thus, our molecular phylogenetic results, haplotype network, and species delimitation analysis, when considered all together, support current *Hermissenda* species limits and relationships and our conclusion that the *Hermissenda* on vessel BF-356 likely did not raft from Japan.

In regard to morphology, the original photographs of the BF-356 *Hermissenda* show white lines on the cerata of some of the nudibranchs, consistent with the external coloration of *H. crassicornis*. Because we only sampled one specimen each of *H. crassicornis* and *H. opalescens* for SEM imaging, we cannot draw any solid conclusions from those data. *Hermissenda crassicornis* having five denticles coming from the central cusp and *H. opalescens* having four denticles may demonstrate characteristic differences between the two species. However, while the numbers of denticles are distinct for each species, they are inconsistent with the characterization of each species according to Lindsay and Valdés (2016); *Hermissenda opalescens* more often have more denticles than *H. crassicornis*. More sampling is required to determine whether denticle number is a reliable character to differentiate between the two species.

Our GBIF map, including iNaturalist data, agrees with the Merlo et al. (2018) range extension of *Hermissenda opalescens* north to Vancouver Island. Our map also shows a more southern extension of *H. crassicornis* than previously understood, extending the area of range overlap between *H. opalescens* and *H. crassicornis*. We note that we used Research Grade observations from iNaturalist, which are fairly reliable, but potential misidentifications may have occurred especially considering the morphological ambiguity between the *Hermissenda* species. Depending on how long this area of range overlap has occurred, *H. opalescens* could potentially be moving north in response to warming ocean temperatures.

Regarding the age of the nudibranchs, lab studies of reared nudibranchs found that largest individuals of *H. crassicornis* from the Monterey Bay grew at a rate of 0.68 mm per day post settlement with about a 35 day veliger stage (Harrigan & Alkon, 1978). According to that rate, the 2.5-cm *Hermissenda* on vessel BF-356, would be approximately 72 days post-hatching. Generation time egg to egg may be as short as 2.5 months (Harrigan & Alkon, 1978), suggesting that if these nudibranchs’ ancestors did raft from Japan, there would have been many generations between the March 2011 earthquake, when the vessel likely departed Japan, and April 2015, when the vessel was found.

While dietary strategies were not the focus of this study, we note that the presence of certain food may have had a role in the presence of *H. opalescens* and *H. crassicornis* on vessel BF-356. While *Hermissenda crassicornis* prefers indigenous food sources to non-indigenous food sources (Kincaid & de Rivera, 2021), it is possible that the local *Hermissenda* larvae in Oregon settled on vessel BF-356 at the cue of the non-indigenous Japanese food sources found on the vessel. Species of *Hermissenda* are known to eat many sessile invertebrates including tunicates, hydroids, and anemones, and specifically *Hermissenda* is known to settle in the presence of and feed on hydroids in the genus *Obelia* which has members native to both Japan and North America (MacDonald & Nybaaken, 1997, Avila, 1998, Megina et al., 2007). *Obelia longissima* is a known prey item of *H. crassicornis* was found on vessel BF-356 (MacDonald and Nybaaken, 1997, Choong et al., 2018) and could have potentially served as a food source and settling cue to both *H. crassicornis* and *H. opalescens*; chemicals associated with nudibranch food are often cues for planktonic nudibranch larvae to settle (Slattery, 1997). One possible scenario is that the planktonic larvae from Oregon settled and were cued to metamorphize by the hydroid (or other) prey found on vessel BF-356. Further work can be done to determine if *Hermissenda* in the eastern Pacific can eat the same prey as *H. emurai* in the western Pacific or cue settling of *H. crassicornis* and *H. opalescens* larvae.

All of our evidence indicates that local *Hermissenda crassicornis* and *H. opalescens* settled on vessel BF-356 after it arrived in the northeast Pacific, rather than native *Hermissenda* from Japan rafting across the Pacific Ocean following the 2011 earthquake and tsunami. While the vessel was intercepted drifting off the central Oregon coast, we do not know its history prior to detection. Large rafted JTMD objects were detected to be drifting along the Oregon and Washington coasts, after acquisition east of coastal boundary currents, both from the north and the south (JTC, personal observations). Given the estimated age of the nudibranchs (above), BF-356 may have been drifting south from, for example, Alaska and/or British Columbia, for some weeks. However, we recognize that such a journey is not impossible, especially if they had a food source like many of the other organisms in the Carlton et al. (2017) study did. Future research on the choosiness in food habits of *Hermissenda* would be impactful in learning about their potential for transoceanic dispersal via a raft that hosted other organisms potentially available for consumption (Carlton et al., 2017).

While our taxonomic conclusions are compelling, we note that the discovery of adult nudibranchs on Japanese tsunami marine debris presents a striking conundrum when considering all of the other 633 (of 634) biofouled sampled JTMD objects analyzed. No other JTMD object studied between 2012 and 2017 supported adult invertebrates from the Northeast Pacific. Instead, recruits of native Northeast Pacifc species consisted, without exception, of 1-2 mm nepionic barnacles (such as *Pollicipes polymerus* and *Balanus glandula)* and nepionic bivalves (such as *Crassadoma gigantea* and *Kellia suborbicularis*). The large adult nudibranchs from JTMD-BF-356 were found inside the vessel, not on the hull where they could have been washed away during transit, suggesting that multiple populations could have been maintained within the vessel over time. If so, and if *Hermissenda* could survive as adults and maintain a population for 1 to 2 years (the estimated departure time of the vessel from Japan), an alternative, but perhaps less likely, hypothesis is that *H. crassicornis* and *H. opalescens* may, in fac,t be in Japan and remain undetected, suggesting that *Hermissenda* ranges in the North Pacific Ocean may need to be investigated further.

## Conclusions

Only Eastern Pacific species, *Hermissenda crassicornis* and *H. opalescens*, were found on vessel BF-356. Our results preserve the currently understood species limits and evolutionary relationships of *Hermissenda* species. The addition of these data also supports the more northern range of *H. opalescens* and expands the southern range of *H. crassicornis*, thus expanding the area of overlap between the species. **While many of the species studied on this floating debris are indicative of transport of northwestern Pacific species to the eastern Pacific, the nudibranchs studied here likely represent secondary colonization of substrate by indigenous species, further complicating the history of this case of invasion.**

## Acknowledgements

We would like to thank L. Esposito for their mentorship of author K. Montana. A. Lam provided training and insights into working in the laboratory and working with haplotype networks. Support in coding for R was provided by G. Rapacciuolo, R. Tarvin, and S. Jacobs. A. Young and A. Miller offered lab moral support. B. Cruz also helped in the lab. We thank S. Rumrill for collecting the nudibranchs from JTMD-BF-356, J. Chapman for photographs of the vessel and the nudibranchs, and G. McDonald and J. Miller for early identification assistance and for further information respectively. We are grateful to N. Treneman for collecting fresh material of Hermissenda for us in 2019 from the Oregon coast. C. Piotrowski L. Kools, and J. Loacker managed our samples in the invertebrate zoology collections at the California Academy of Sciences. This project originated as a National Science Foundation Research Experience for Undergraduates project, so we thank the NSF for the grant DBI-1852276 to L. Esposito and R. Johnson that supported this work. We also thank the fellow interns in the Summer Systematics Institute REU program for community and support throughout the project. We thank the nudibranchs for giving their lives to science. We thank the Ramaytush Ohlone and the Confederated Tribes of Siletz Indians for stewarding the land and coasts where our work took place.

## Works Cited

Arce-Valdés LR, Sánchez-Guillén RA. The evolutionary outcomes of climate-change-induced hybridization in insect populations. Curr Opin Insect Sci. 2022 Dec 1;54:100966. doi: 0.1016/j.cois.2022.100966.

Avila C. Competence and metamorphosis in the long-term planktotrophic larvae of the nudibranch mollusc Hermissenda crassicornis (Eschscholtz, 1831). Journal of Experimental Marine Biology and Ecology. 1998: 231(1)81–117.10.1016/S0022-0981(98)00093-8.

Baba K. Opisthobranchia of Japan (II). J Dep Agric. 1937;5(7):289–344.

Barnard KH. South African nudibranch *Mollusca*, with descriptions of new species, and a note on some specimens from Tristan d’Acunha. Ann S Afr Mus. 1927;25:171–215. https://www.biodiversitylibrary.org/item/126046.

Behrens DW, Fletcher K, Hermosillo A, Jensen GC. (2022). Nudibranchs & sea slugs of the Eastern Pacific. Mola Marine. 2022;170.

Bergh R. Beiträge zur kenntniss der aeolidiaden. VI. Verh Zool Bot Ges Wien. 1878;28: 553–584.

Burghardt I, Wägele H. A new solar powered species of the genus *Phyllodesmium* Ehrenberg, 1831 (Mollusca: Nudibranchia: Aeolidoidea) from Indonesia with analysis of its photosynthetic activity and notes on biology. Zootaxa. 2004;596(1): 1–18. doi: 10.11646/zootaxa.596.1.1.

Carlton JT, Chapman JW, Geller JB, Miller JA, Carlton DA, McCuller MI, et al. Tsunami-driven rafting: Transoceanic species dispersal and implications for marine biogeography. Science. 2017;357(6358):1402–1406. doi: 10.1126/science.aao1498.

Carlton JT, Geller JB. Ecological roulette: The global transport of nonindigenous marine organisms. Science. 1993 July 2;261(5117):78–82. doi: 10.1126/science.261.5117.78.

Carlton JT, Chapman JW, Geller JB, Miller JA, Ruiz GM, Carlton DA, et al. Ecological and biological studies of ocean rafting: Japanese tsunami marine debris in North America and the Hawaiian islands. Aquat Invasions. 2018 Feb;13(1):1–9. doi: 10.3391/ai.2018.13.1.01.

Chernomor O, von Haeseler A, Minh BQ. Terrace aware data structure for phylogenomic inference from supermatrices. Syst Biol. 2016;65: 997–1008. doi: 10.1093/sysbio/syw037.

Choong HHC, Calder DR, Chapman JW, Miller JA, Geller JB, Carlton JT. Hydroids (Cnidaria: Hydrozoa: Leptothecata and Limnomedusae) on 2011 Japanese tsunami marine debris landing in North America and Hawai‘i, with revisory notes on *Hydrodendron* Hincks, 1874 and a diagnosis of Plumaleciidae, new family. Aquat Invasions. 2018 Feb;13(1):43–70. doi: 10.3391/ai.2018.13.1.01.

Churchill CKC, Alejandrino A, Valdés A, Foighil DO. Parallel changes in genital morphology delineate cryptic diversification of planktonic nudibranchs. Proc Biol Sci. 2013 Aug 22;280(1765): 20131224. doi: 10.1098/rspb.2013.1224.

Cooper JG. On new or rare Mollusca inhabiting the coast of California. No. II. Proc Calif Acad Sci. 1863;3:56–60.

Cooper JG. Some new genera and species of California Mollusca. Proc Cal Acad Nat Sci. 1862;2:202–205, 207.

Craig MT, Burke J, Clifford K, Mochon-Collura E, Chapman JW, Hyde JR. Trans-Pacific rafting in tsunami associated debris by the Japanese yellowtail jack, *Seriola aureovittata* Temminck & Schlegel, 1845 (Pisces, Carangidae). Aquatic Invasions 2018 13(1):173–177

Engel H. Westindische opisthobranchiate Mollusken. Bijdragen tot de kennis der fauna van Curaçao. Resultaten eener reis van Dr. C. J. van der Horst in 1920. Contrib Zool. 1925;24: 33–80.

Eschscholtz JF. Zoologischer atlas—Beschreibungen neuer thierarten, während des Flottcapitains von Kotzebue zweiter reise um die welt, auf der Russisch-Kaiserlichen kriegsschlupp Predpriaetië in den jahren 1823–1826. G. Reimer, Berlin. 1831;4:1–19.

Folmer O, Black M, Hoeh W, Lutz R, Vrijenhoek R. DNA primers for amplification of mitochondrial cytochrome c oxidase subunit I from diverse metazoan invertebrates. Mol Mar Biol Biotechnol. 1994 Oct;3(5)294–299.

Garroway CJ, Bowman J, Cascaden TJ, Holloway GL, Mahan CG, Malcolm JR, et al. Climate change induced hybridization in flying squirrels. Glob Chang Biol. 2010;16(1):113–121. doi: 10.1111/j.1365-2486.2009.01948.x.

GBIF.org. GBIF Occurrence Download; 2022 [cited 2022 Jul 6]. Database: GBIF [Internet]. Available from: 10.15468/dl.h4u2kp.

GBIF.org. GBIF Occurrence Download; 2022 [cited 2022 Jul 6]. Database: GBIF [Internet]. Available from: 10.15468/dl.v3ahgx

GBIF.org. GBIF Occurrence Download; 2022 [cited 2022 Dec 14]. Database: GBIF [Internet]. Available from: 10.15468/dl.csmfq6

GBIF.org. GBIF Occurrence Download; 2022 [cited 2022 Dec 14]. Database: GBIF [Internet]. Available from: 10.15468/dl.fbhsmh

Gollasch, S. Is ballast water a major dispersal mechanism for marine organisms? In: Wolfgang N, editor. Biological Invasions. Berlin, Heidelberg: Springer; 2007. pp. 49–57. 10.1007/978-3-540-36920-2_4.

Goodheart JA, Bazinet AL, Valdés Á, Collins AG, Cummings MP. Prey preference follows phylogeny: evolutionary dietary patterns within the marine gastropod group Cladobranchia (Gastropoda: Heterobranchia: Nudibranchia). BMC Evol Biol. 2017 Oct 26;17(1): 221. doi: 10.1186/s12862-017-1066-0.

Goodheart JA, Bleidißel S, Schillo D, Strong EE, Ayres DL, Preisfeld A, Collins AG, Cummings MP, Wägele H. Comparative morphology and evolution of the cnidosac in Cladobranchia (Gastropoda: Heterobranchia: Nudibranchia). Front Zool. 2018;15 43. doi: 10.1186/s12983-018-0289-2.

Hadfield MG. The biology of nudibranch larvae. Oikos. 1963;14(1):85–95. doi: 10.2307/3564960.

Harrigan JF, Alkon DL. Larval rearing, metamorphosis, growth and reproduction of the eolid nudibranch *Hermissenda crassicornis* (Eschscholtz, 1831) (Gastropoda: Opisthobranchia). The Biological Bulletin. 1978 Jun;154(3):430–9.

Hart LC, Goodman MC, Walter RK, Rogers-Bennett L, Shum P, Garrett AD, Watanabe JM, O’Leary JK. Abalone recruitment in low-density and aggregated populations facing climatic stress. J Shellfish Res. 2020 Aug;39(2): 359–373. doi:10.2983/035.039.0218.

Highsmith RC. Floating and algal rafting as potential dispersal mechanisms in brooding invertebrates. Mar. Ecol. Prog. Ser. 1985;25:169–179.

Hoang DT, Chernomor O, von Haeseler A, Minh BQ, Vinh LS.UFBoot2: Improving the ultrafast bootstrap approximation. Mol Biol Evol. 2018;35: 518–522. doi: 10.1093/molbev/msx281.

Hobbs JA, Frisch Aj, Allen GR, Van Herwerden L. Marine hybrid hotspot at Indo-Pacific biogeographic border. Biol Lett. 2009 Apr 23;5(2):258–261. doi: 10.1098/rsbl.2008.0561.

Kalyaanamoorthy S, Minh BQ, Wong TKF, von Haeseler A, Jermiin LS. ModelFinder: fast model selection for accurate phylogenetic estimates. Nat Methods. 2017;14: 587–589. doi: 10.1038/nmeth.4285.

Kincaid ES, de Rivera CE. Predators associated with marinas consume indigenous over non-indigenous ascidians. Estuaries Coast. 2021 May 1;44(3):579–588. doi: 10.1007/s12237-020-00793-2.

Leigh JW, Bryant D. popart: full-feature software for haplotype network construction. Methods Ecol Evol. 2015;6:1110–1116. doi: 0.1111/2041-210X.12410.

Lindsay T, Valdés Á. The model organism *Hermissenda crassicornis* (Gastropoda: Heterobranchia) is a species complex. PLoS One. 2016 Apr 22;11(4):e0154265. doi: 10.1371/journal.pone.0154265.

Madeira F, Pearce M, Tivey ARN, Basuktar P, Lee J, Edbali O, et al. Search and sequence analysis tools services from EMBL-EBI in 2022. Nucleic Acids Res. 2022 Apr 12;50(W1):W276–W279. doi: 10.1093/nar/gkac240.

MacDonald G, Nybaaken J. 1997. List of the Worldwide Food Habits of Nudibranchs. https://escholarship.org/uc/item/0g75h1q3

Marcus E. On Opisthobranchia from Brazil (2). Zool J Linn Soc. 1957;43(292): 390–486. doi: 10.1111/j.1096-3642.1957.tb01559.x.

McDonald GR. A review of the nudibranchs of the California coast. Malacologia. 1983;24:114–276.

McDonald GR, Nybakken JW. List of the worldwide food habits of nudibranchs. UC Santa Cruz Previously Published Works. 1997.

Megina C, Cervera JL. Diet, prey selection and cannibalism in the hunter opisthobranch *Roboastra europaea*. J Mar Biol Assoc UK. 2003 June;83(3):489–95. doi: 10.1017/S0025315403007392h.

Merlo EM, Milligan KA, Sheets NB, Neufeld CJ, Eastham TM, Estores-Pacheco ALK, et al. Range extension for the region of sympatry between the nudibranchs *Hermissenda opalescens* and *Hermissenda crassicornis* in the northeastern Pacific. FACETS. 2018 Oct 1;3(1):764–776. doi: 10.1139/facets-2017-0060.

Miller MC. Aeolid nudibranchs (Gastropoda: Opisthobranchia) of the family Glaucidae from New Zealand waters. Zool J Linn Soc. 1974;54:31–61.

Moore EJ, Gosliner TM. Molecular phylogeny and evolution of symbiosis in a clade of Indopacific nudibranchs. Mol Phylogenet Evol. 2011 Jan;58(1): 116–123. doi: 10.1016/j.ympev.2010.11.008.

Nguyen LT, Schmidt HA, von Haeseler A, Minh BQ. IQ-TREE: A fast and effective stochastic algorithm for estimating maximum likelihood phylogenies. Mol Biol Evol. 2015;32: 268–274. Doi: 10.1093/molbev/msu300.

O’Donoghue CH. Notes on the taxonomy of nudibranchiate Mollusca from the Pacific coast of North America. I. On the identification of Cavolina (i.e. Hermissenda) crassicornis of Eschscholtz. Nautilus. 1922; 35:74–77.

Pebesma E. Simple features for R: Standardized support for spatial vector data. The R Journal 10. 2018;1:439–446. doi: 10.32614/RJ-2018-009.

Pola M, Gosliner TM. The first molecular phylogeny of cladobranchian opisthobranchs (Mollusca, Gastropoda, Nudibranchia). Mol Phylogenet Ev. 2010 Sep;56(3): 931–941. doi: 10.1016/j.ympev.2010.05.003.

Puillandre N, Lambert A, Brouillet S, Achaz G. ABGD, Automatic Barcode Gap Discovery for primary species delimitation. Mol Ecol. 2012 Apr;21(8):1864–77. doi: 10.1111/j.1365-294X.2011.05239.x. https://bioinfo.mnhn.fr/abi/public/abgd/abgdweb.html

Ronquist F, Teslenko M, van der Mark P, Ayres DL, Darling A, Höhna S, et al. MRBAYES 3.2: Efficient Bayesian phylogenetic inference and model selection across a large model space. Syst. Biol. 2012;61:539–542. doi: 10.1093/sysbio/sys029.

Shields C. Nudibranchs of the Ross Sea, Antarctica: phylogeny, diversity, and divergence. Master’s thesis. 2009 Aug 1. https://tigerprints.clemson.edu/all_theses/637.

Simkanin C, Carlton JT, Steves B, Fofonoff P, Nelson JC, Clarke Murray C, Ruiz GM. Exploring potential establishment of marine rafting species after transoceanic long-distance dispersal. Glob Ecol Biogeogr. 2019;28(5):588–600. doi: 10.1111/geb.12878.

Simkanin C, Davidson IC, Therriault TW, Jamieson G, Dower JF. Manipulating propagule pressure to test the invasibility of subtidal marine habitats. Biol Invasions. 2017 May 1;19:1565–1575. doi: 10.1007/s10530-017-1379-3.

Slattery M. Chemical cues in marine invertebrate larval settlement. Fouling Organisms of the Indian Ocean. CRC Press. 1997.

Thiel M, Gutow L. The ecology of rafting in the marine environment. I. The floating substrata. In: Oceanography and marine biology: An annual review. 2005;42:181–264.

Trickey JS, Thiel M, Waters JM. Transoceanic dispersal and cryptic diversity in a cosmopolitan rafting nudibranch. Invertebr Syst. 2016 June;30(3): 290–301. doi: 10.1071/IS15052.

Trifinopoulos J, Nguyen LT, von Haeseler A, Minh BQ. W-IQ-TREE: A fast online phylogenetic tool for maximum likelihood analysis. Nucleic Acids Res. 2016 Jul 8; 44(W1):W232–235. doi: 10.1093/nar/gkw256.

Turgeon DD. Common and scientific names of aquatic invertebrates from the United States and Canada: Mollusks. 1st ed. American Fisheries Society; 1988.

